# Unveiling immune targets via TriStageHLA: high-throughput screening of peptide presentation, peptide–HLA binding, and immunogenicity

**DOI:** 10.64898/2026.10.29.762831

**Authors:** Tianchi Lu, Zikun Wang, Yuchen Wang, Zeji Li, Tianhang Chen, Ling Cho, Zhongyu Yao, Xiangtao Li, Ka-Chun Wong

## Abstract

High-throughput screening of HLA class I epitopes underpins neoantigen vaccine design, T cell therapies, and pathogen immune monitoring, yet proteome-scale prioritization remains challenging under severe class imbalance. We present TriStageHLA, a compact three-stage framework that decomposes epitope discovery into peptide-only screening (PRE), allele-specific binding/presentation prediction (BIND), and immunogenicity prediction (IM). PRE performs early pruning via a LoRA-adapted ESM-2 classifier; BIND integrates quantitative binding affinity and eluted ligand data with a dual-branch architecture combining global cross-attention and local motif extraction; IM is fine-tuned on curated human T-cell assay data with a decoupled LoRA head. Trained on IEDB-derived datasets with per-allele balancing and allele-discordant supervision, TriStageHLA-BIND achieves AUROC of 96.2% and AUPRC of 95.2% on combined BA/EL evaluation, while TriStageHLA-IM reaches AUROC of 81.4% and AUPRC of 73.8% on held-out immunogenicity data, improving over the strongest immunogenicity baselines ImmuneApp (70.7% / 63.6%) and BigMHC-IM (60.1% / 51.5%). With substantially fewer trainable parameters than prior models, TriStageHLA enables efficient, calibrated epitope prioritization across binding, presentation, and immunogenicity. Code and pretrained weights are available at https://github.com/SkywalkerLuke/TriStageHLA.

## 1 Introduction

Cytotoxic CD8+ T cells maintain immune surveillance by recognizing short peptides displayed by HLA class I molecules, eliminating infected or malignant cells and shaping adaptive immunity [1–6]. This gatekeeping step underpins diverse applications—from peptide vaccines for emerging pathogens and individualized T cell therapies to neoantigen discovery, autoimmunity research, and transplant immunology [7–12]. Despite substantial advances in LC–MS/MS immunopeptidomics and binding/stability assays, truly immunogenic peptide–HLA (pHLA) ligands are rare [13–15]. Comprehensive experimental discovery at proteome scale remains resource-intensive, motivating computational strategies that can prioritize candidates while preserving decision quality under pronounced class imbalance [13, 14, 16, 17].

From a translational standpoint, the practical bottleneck is the mismatch between what each experimental assay measures and what a screen ultimately needs. Binding assays quantify how tightly a peptide attaches to an HLA molecule, providing precise, allele-specific labels, but they are laborious and therefore sparse and skewed toward a handful of well-studied HLA alleles [18, 19]. Eluted-ligand (EL) experiments, which recover peptides actually displayed on the cell surface, scale far better and give unambiguous HLA restriction in mono-allelic systems [20, 21]. Yet because EL data reveal what is presented but not what is absent, models trained on them are typically forced to inject large numbers of random “decoy” negatives; this shortcut inflates apparent accuracy and biases predictions toward the abundant, easy-to-reject background [15, 22]. In a clinical screen, where a single tumor exome yields tens of thousands of candidate peptides but only a handful are therapeutically relevant, that bias is precisely what must be avoided: dismissing a rare true epitope is far costlier than testing a few extra candidates. These trade-offs motivate separating what each kind of evidence can reliably teach—broad peptide enrichment versus allele-specific binding—and aligning the training data to each decision, as elaborated below.

Computationally, antigen prediction comprises related but distinct decisions: whether a peptide exhibits epitope-like sequence features; whether it binds a particular HLA allele; and whether it elicits T cell responses in humans. A peptide-only presentation classifier can front-load candidate pruning, and allele-agnostic presentation modeling has been shown to serve as an effective front-end filter at scale [23].

In practice, however, earlier implementations were heavyweight (on the order of hundreds of millions of parameters) and did not support parameter-efficient adaptation of the learned representations for downstream tasks such as allele-specific binding or immunogenicity prediction. These limitations, together with the complementary strengths and weaknesses of BA and EL supervision outlined above, motivate the compact, modular redesign presented in the next section, where we detail how presentation enrichment, allele-specific binding, and immunogenicity are trained and coupled.

To that end, we introduce TriStageHLA, a compact framework that separates epitope discovery into three intuitive questions, asked in sequence (Figure 1). The first stage asks only whether a peptide looks like something the immune system will display, without committing to any particular HLA allele; it rapidly discards most of the proteome while keeping the rare true epitopes, and it does so without the large random-negative ratios that bias conventional screens [13, 24, 25]. The second stage asks which HLA allele a peptide binds, learning precision from quantitative binding measurements and from cases where the same peptide binds one allele but not another, with balanced positive and negative examples per allele [21, 26–30]. The third stage asks, separately, whether a peptide will actually trigger a T-cell response, trained on carefully curated human T-cell assay data under strict quality filters [31– 34]; the curation, filtering, and partitioning procedures are detailed in the Methods (Datasets) section. We deliberately treat immunogenicity as a separate module with its own curated data, rather than as the ultimate endpoint, so that binding and T-cell responsiveness are not conflated.

**Fig. 1.**
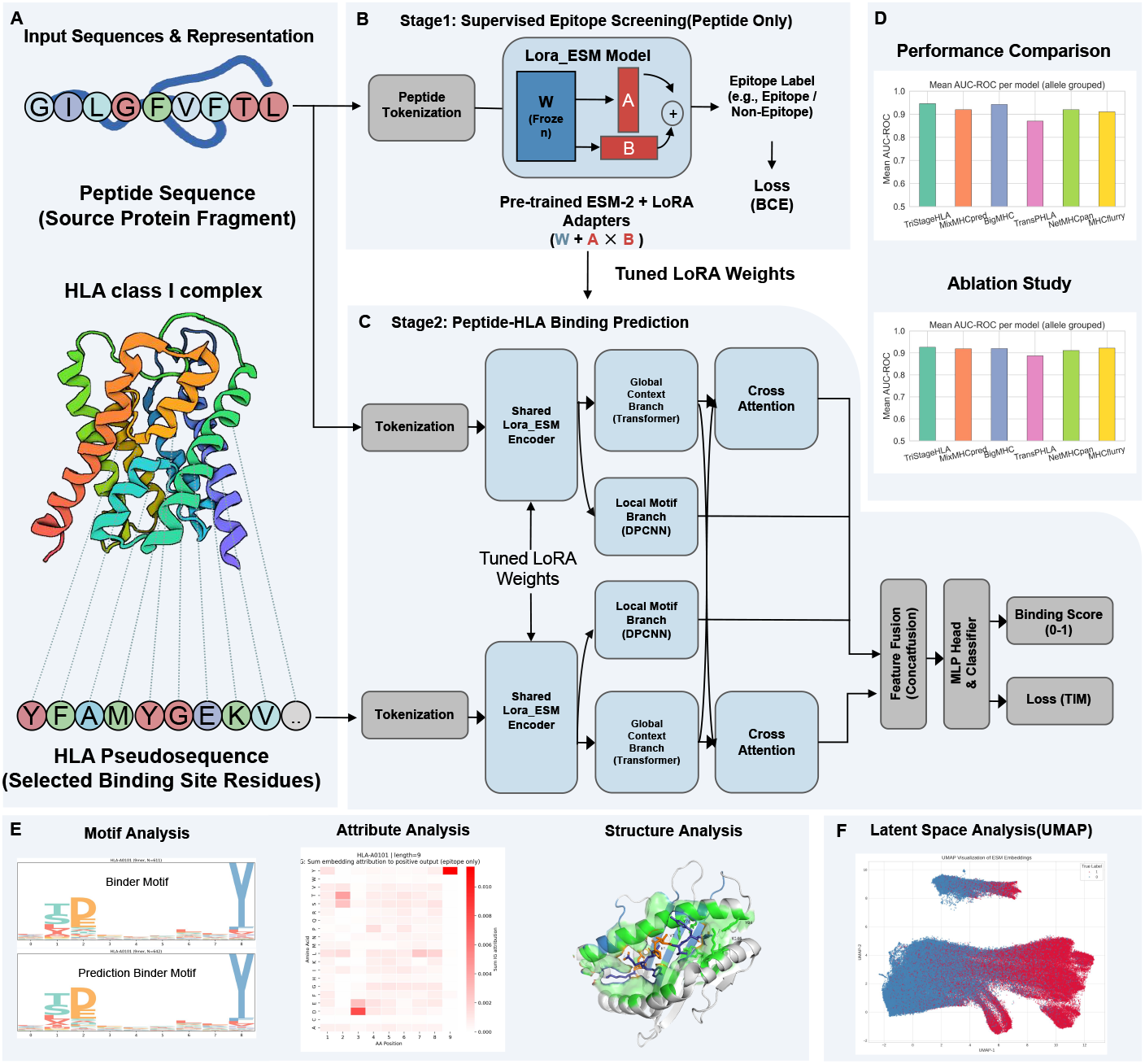
Overview of the peptide–HLA binding prediction framework. (A) Inputs include peptide sequences and HLA pseudosequences, with a schematic of their complex. (B) Stage 1: Pretraining with LoRA-tuned ESM2 to learn protein language representations. (C) Stage 2: Binding prediction with a dual-branch architecture—global-context modeling (Transformer + cross-attention) and local-motif extraction (CNN)—whose fused outputs yield the final classification. (D) Performance comparison and ablation study showing the full model outperforming ablated variants and prior SOTA. (E) Interpretability analyses, including sequence motif discovery, attribution maps, and structure-based interaction analysis. (F) UMAP embeddings illustrating separation of binders vs. non-binders.

Across independent mono-allelic benchmarks, this design yields presentation-oriented performance with a substantially reduced trainable footprint, enabling early pruning that lowers the end-to-end runtime for proteome-scale scans without compromising recall [15, 20]. On allele-specific binding, TriStageHLA performs competitively with or exceeds established predictors, benefiting from quantitative IC50 supervision and peptide-wise allele discordance; on immunogenicity, it improves precision at practical decision thresholds while maintaining calibrated operating points [34– 36]. Concretely, TriStageHLA-BIND attains AUROC 96.2% and AUPRC 95.2% on BA/EL evaluation and leads at the allele level (AUROC 94.5%, AUPRC 89.7%, ACC 88.8%, MCC 0.717), with BigMHC [34] and MixMHCpred3.0 as the second- and third-best baselines; by length, it preserves discrimination for longer peptides (e.g., 13–14-mers AUROC 92–94%, AUPRC 76–86%) [34, 36, 37]. On the immunogenicity task, TriStageHLA-IM achieves AUROC 81.4% and AUPRC 73.8% on held-out data, outperforming the strongest immunogenicity baselines ImmuneApp (70.7% / 63.6%) and BigMHC-IM (60.1% / 51.5%). End-to-end, the modular pipeline (TriStageHLA-PRE → TriStageHLA-BIND → TriStageHLA-IM) supports aggressive yet safe early pruning and improves precision at practical immunogenicity thresholds while maintaining calibration and recall. In practical terms, this yields a candidate list that is both shorter and better focused—fewer peptides to validate experimentally while retaining the therapeutically relevant ones—directly reducing the cost and turnaround of personalized epitope discovery.

Model interpretability further supports the structural and biochemical validity of the predictions. Attribution analyses and sequence logos recover canonical anchor positions and residue preferences, and structure-aware examinations align learned importance with peptide–HLA contact residues and binding pockets [38–40]. Over-all, these results suggest that the proposed modular framework—with peptide-first screening, allele-specific binding, and curated immunogenicity—offers a practical and interpretable approach for immunopeptidomics across infectious disease, oncology, autoimmunity, and transplantation [13, 16].

## 2 Results

### 2.1 Comparative results for peptide-only screening and allele-aware binding/presentation

#### 2.1.1 Peptide-only presentation screening

TriStageHLA-PRE reduces the trainable parameter count by approximately one order of magnitude relative to TransHLA (about 35M versus *∼* 0.7B parameters) by employing LoRA (Low-Rank Adaptation)-based [41] fine-tuning, which updates a small set of low-rank adapters instead of the full model. Under the same protocol with metrics computed from raw logits, it preserves or slightly improves discrimination on the held-out evaluation: as summarized in Figure 2A, accuracy reached 84.85% and AUROC reached 92.05%, marginally exceeding TransHLA’s 84.70% and 91.90%. The LoRA adapters capture generalizable peptide features and retain weights that serve as an initialization for subsequent, task-specific fine-tuning stages in the pipeline.

**Fig. 2.**
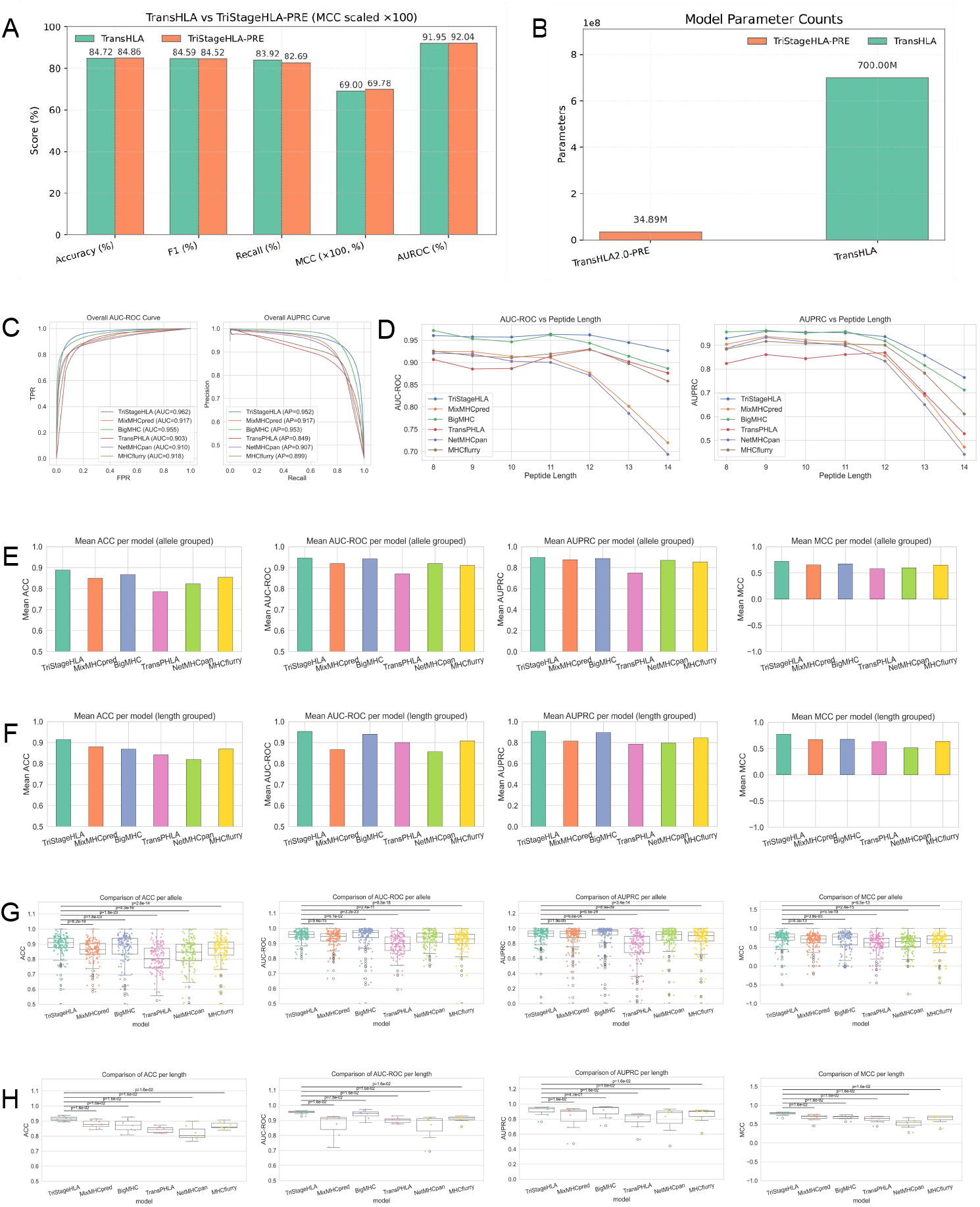
Independent tests on presentation and bind with stratified analyses. (A–B) Performance versus capacity, comparing TriStageHLA-PRE and TransHLA across standard metrics; parameter counts shown for both models (*≈* 35M frozen encoder + LoRA vs 0.7B). (C) Overall performance using raw logits, aggregated across alleles (AUROC, AUPRC, accuracy). (D) Length-wise performance (8–14-mers) across peptide lengths. (E) Allele-wise mean bars. (F) Length-wise mean bars. (G) Allele-wise box-and-scatter with paired bootstrap/DeLong tests versus the strongest baseline (BH-adjusted). (H) Length-wise box-and-scatter with the same statistical testing.

#### 2.1.2 Allele-aware peptide–HLA binding/presentation

On the combined BA/EL evaluation, TriStageHLA-BIND achieved AUROC 96.2% and AUPRC 95.2%, improving AUROC from 95.5% to 96.2% versus BigMHC (AUPRC comparable) and from 92.7% to 96.2% and 92.7% to 95.2% versus MixMHCpred3.0 [37] (AUROC and AUPRC). Figure 2C summarizes these overall results, with box-and-scatter views in Figure 2F–G supporting reduced dispersion and paired gains.

At the allele level, TriStageHLA-BIND leads on discrimination with AUROC 94.54% and AUPRC 89.74%, alongside balanced accuracy (ACC 88.82%, MCC 71.69%). Among baselines, BigMHC and MixMHCpred3.0 perform best overall (second and third, respectively): BigMHC reaches AUROC 94.21% and AUPRC 88.77%, while MixMHCpred3.0 reaches AUROC 91.94% and AUPRC 87.46%. Figure 2E shows allele-stratified bars, where a majority of alleles favor TriStageHLA-BIND over the next-best baselines; Figure 2F–G indicate reduced variance and paired improvements. Detailed per-allele results are provided in Supplementary Table 1.

Because the clinical question is whether TriStageHLA-BIND helps precisely where established tools are weakest, we stratified the per-allele results by difficulty along two axes: sample scarcity (“rare” alleles with *<*200 training instances) and binding-site distance to the nearest training allele (“distant” alleles with pseudo-sequence distance *>*0.1, as defined in the distance-based evaluation below). The gain over BigMHC grows monotonically with difficulty: on rare alleles (*n* = 34) TriStageHLA-BIND attains the highest mean AUROC of all six methods (0.899) and significantly outperforms BigMHC (0.857; +0.042, *p* = 0.019); on the hardest rare-and-distant intersection (*n* = 10) the advantage is largest (+0.080 AUROC, +0.172 AUPRC). The only group where BigMHC edges ahead is the distant-but-not-rare subset (*n* = 11), i.e., alleles that are structurally atypical yet still training-rich, where BigMHC’s larger capacity holds a narrow lead. This monotonic pattern—greater difficulty, greater TriStageHLA-BIND advantage—directly addresses the reviewer’s question of whether the model adds value specifically on rare and under-represented alleles. The overlap between the rare and distant subsets and the per-group comparisons are provided in Supplementary Figure 10.

As stratified into different k-mers, TriStageHLA-BIND sustains stronger performance for longer peptides. For 13-mers, AUROC increases from 91.39% to 94.47% versus BigMHC and from 80.16% to 94.47% versus MixMHCpred3.0 (AUPRC from 81.59% to 85.75% and from 68.85% to 85.75%, respectively). For 14-mers, AUROC increases from 88.65% to 92.67% versus BigMHC and from 71.99% to 92.67% versus MixMHCpred3.0 (AUPRC from 71.23% to 76.49% and from 47.18% to 76.49%). Figure 2D visualizes these trends, highlighting robust performance at 13–14 residues relative to steeper declines for other methods, while BigMHC and MixMHCpred3.0 remained as the strongest baselines. Detailed k-mer–stratified metrics are provided in Supplementary Table 2.

### 2.2 Interpretability and motif validation

We selected HLA-A*02:01 as a well-characterized benchmark allele with abundant training data, and HLA-B*57:03 and HLA-C*12:02 as representatives of less frequently profiled HLA-B and HLA-C alleles, to illustrate performance on both well-resourced and harder cases. Using the three representative alleles shown in Figure 3 as prototypes for HLA-A, HLA-B, and HLA-C, we performed a stepwise interpretability analysis spanning sequence motifs and peptide-length preferences. We first examined the sequence logos in Figure 3A–C, where the left panels summarize motifs derived from experimentally observed ligands and the right panels summarize motifs from the model’s top-scoring, length-matched, non-redundant peptides. Across all three alleles, the predicted logos closely mirror the true motifs at the key high-information positions, indicating that the model recovers allele-specific binding rules. Detailed sequence logos for all alleles, including predicted logos, are provided in Supplementary Figures 4–6. For the HLA-A representative HLA-A*02:01 (Figure 3A), TriStageHLA-BIND accurately reproduces the canonical dual-anchor pattern, highlighting a strong preference for a hydrophobic leucine at P2 and valine/leucine at the C-terminal anchor (PΩ; for 9-mers this corresponds to P9). For the HLA-B representative HLA-B*57:03 (Figure 3B), the predicted motif captures enriched polar residues (notably S/T) at motif-defining N-terminal positions together with a dominant aromatic/hydrophobic signature (F/W/L) at the C-terminal anchor, consistent with the allele’s characteristic “polar-front / hydrophobic-tail” binding mode. For the HLA-C representative HLA-C*12:02 (Figure 3C), the model likewise recapitulates the true motif, emphasizing an N-terminal preference for A and marked enrichment of Y/F and L at key positions, demonstrating robust motif learning in HLA-C alleles as well.

**Fig. 3.**
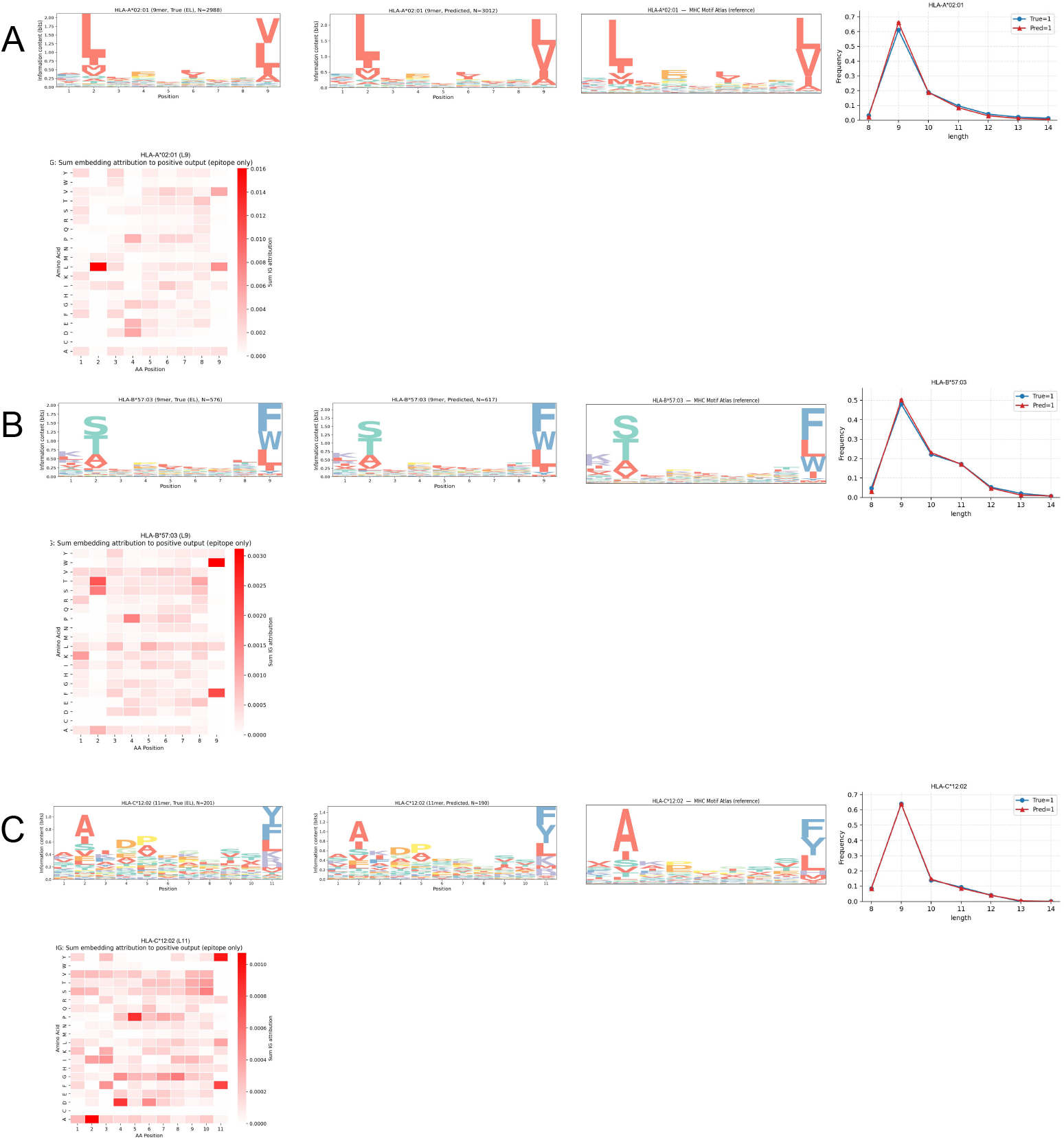
Interpretability across alleles via sequence motifs, attributions, and motif references. (A–C) Three representative alleles, each shown as a row of five complementary views: sequence logos of experimentally observed (EL-positive) ligands, sequence logos of the model’s top-scoring peptides, the independently reconstructed reference motif from the MHC Motif Atlas (Tadros et al., 2023), the peptide-length distribution, and the attribution-derived preference map. Anchor positions remain consistent across all views.

To anchor this comparison in an independent, training-free reference, we reconstructed 9-mer position-weight matrices from the MHC Motif Atlas (Tadros et al., 2023) and compared them against the model’s top-1,000 predicted peptides per allele. The predicted motifs are highly concordant with the Atlas reference (mean Jensen–Shannon divergence 0.026, 0.027, and 0.013 for HLA-A*02:01, HLA-B*57:03, and HLA-C*12:02, respectively), with top-position amino-acid identity of 55.6%, 66.7%, and 66.7%, and the canonical anchor residues recover fully in every case (HLA-A*02:01 P2 L/I/M and P9 L/V/I; HLA-B*57:03 P2 S/T/A and P9 F/L/W; HLA-C*12:02 P2 A/I and P9 Y/F/L). This confirms that the recovered motifs are not an artifact of comparing against the training-derived logos alone.

We next compared peptide-length frequency distributions for each allele (middle plots in Figure 3A–C) using both the training ligands and the model’s high-confidence predictions. All three alleles exhibit a pronounced dominant peak at length 9, whereas other lengths form only minor secondary peaks. This shared pattern indicates that 9-mer peptides represent the major presentation class in these representative alleles. Comprehensive length-distribution plots for all alleles are provided in Supplementary Figures 1–3. Accordingly, we selected the length-9 subset as a unified and statistically well-supported basis for attribution analysis.

Building on this choice, we aggregated token-level attributions over correctly ranked 9-mer peptides to obtain position-specific amino-acid preference maps (right heatmaps in Figure 3A–C). The attribution landscapes align tightly with the motif peaks in each case. In HLA-A*02:01, the strongest attribution weights concentrate at P2 and P9, with maximal contributions assigned specifically to L at P2 and V/L at P9, confirming that the model’s decisions are dominated by the terminal hydrophobic anchors. In HLA-B*57:03, attributions form a clear two-peak structure: elevated contributions at N-terminal motif-defining positions favoring S/T, and the largest weights at the C-terminal anchor favoring F/W/L, reinforcing that the model relies on allele-specific anchor residues rather than diffuse positional signals. In HLA-C*12:02, attribution hotspots again coincide with the high-information logo columns, showing strong dependence on the N-terminal A preference and on Y/F and L at core motif positions. Full attribution heatmaps across alleles are provided in Supplementary Figures 7–9. Together, these results demonstrate that TriStageHLA-BIND’s discriminative cues are sharply localized to motif-defining residues across HLA-A, HLA-B, and HLA-C, providing a coherent sequence-level explanation of its predictions.

Finally, we assessed structure-level validity by projecting attribution weights onto crystallographic peptide–HLA class I complexes for the same alleles (Supplementary Figure 12). For HLA-A*02:01 bound to KLYQNPTTYI (PDB 3MGT), the highest-attribution residues map to the N-terminal P2 anchor engaging the B pocket and the C-terminal P9 anchor engaging the F pocket, which are the densest contact regions in the binding groove. For HLA-B*57:03 bound to KGFNPEVIPMF (PDB 2HJK), attribution hotspots likewise co-localize with the principal N-terminal and C-terminal anchoring regions, while central positions show weaker secondary contributions corresponding to backbone hydrogen bonds and supportive side-chain contacts. For HLA-C*12:02 bound to KAYNVTQAF (PDB 9F13), the mapped attributions again highlight terminal anchors consistent with the inferred motif, confirming that the model’s key sequence signals reflect genuine structural constraints. The strong agreement between motif logos, attribution maps, and physical contact patterns across all three alleles indicates that the model learns meaningful determinants of peptide–HLA presentation rather than spurious correlations.

### 2.3 Generalization across external datasets and out-of-distribution alleles

#### 2.3.1 External dataset generalization (independent HLA–peptide pairs)

We evaluated binding/presentation predictions on an external test set whose positive HLA–peptide pairs were drawn entirely from The Cancer Epitope Database and Analysis Resource (CEDAR) [42] and were not included at any development stage. Negatives were generated by aligning to a non-redundant protein database, randomly extracting peptide segments, and randomly pairing them with HLA alleles; these negatives were likewise excluded from model development. This design enables a direct assessment of out-of-sample generalization.

Length-stratified performance (Figure 4A) shows that TriStageHLA-BIND maintains its advantage across peptide lengths: AUROC remained above 80% for all lengths, with complete superiority over BigMHC at lengths 10–12, and AUPRC comparable to BigMHC across lengths. In aggregate comparisons (Figure 4B), TriStageHLA-BIND exhibited the smallest drop relative to internal validation, achieving AUROC of 94.2% versus 93.0% for MixMHCpred3.0 and 92.0% for BigMHC; AUPRC reached 95.0% versus 94.7% for MixMHCpred3.0. Allele- and length-wise summaries (box–scatter and bar plots; Figure 4C) with two-tailed Wilcoxon signed-rank tests and multiple-testing adjustment indicate stable allele-wise performance with fewer outliers in AUROC compared with MixMHCpred3.0 and BigMHC, and stronger length-wise robustness. Detailed length- and allele-stratified results are provided in Supplementary Tables 3 and 4.

**Fig. 4.**
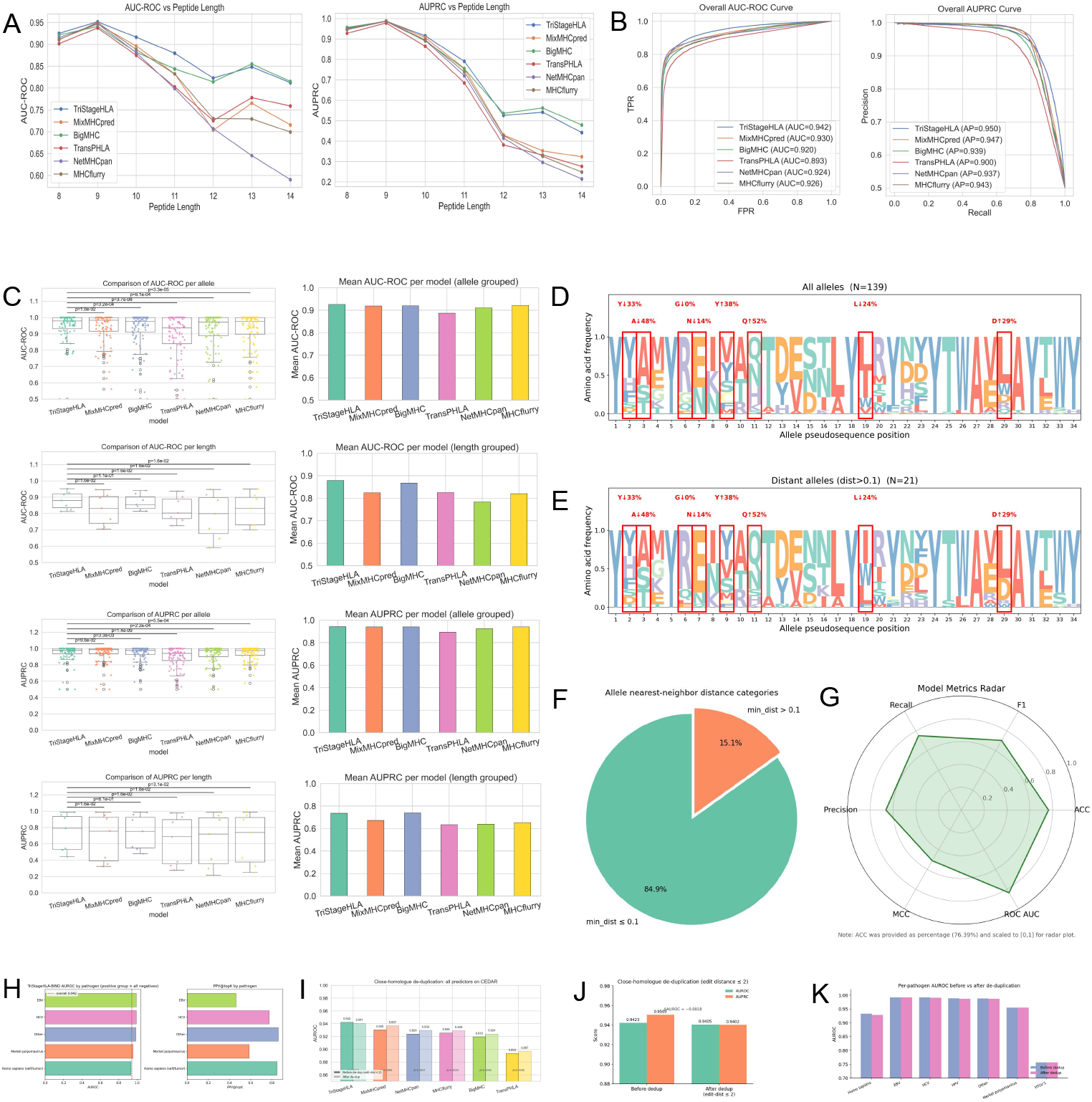
Generalization on CEDAR with allele- and length-stratified views and allele-distance analysis. (A) Length-wise AUROC/AUPRC (8–14-mers) on CEDAR with 95% CIs. (B) Overall AUROC/AUPRC on CEDAR using raw logits. (C) Stratified CEDAR performance: per-allele scatter/line summaries and per-length bar plots with paired tests versus baseline (BH-adjusted). (D) Global sequence logo aggregated over all alleles. (E) Sequence logos for alleles with pseudo-sequence distance *>* 0.1, indicating motif conservation under larger allele shifts. (F) Distribution of pairwise allele distances, showing proportions of near vs. far neighbors. (G) Far-allele stress test: radar plot with the most distant alleles held out for testing and the remainder used for train/validation. (H) AUROC on CEDAR stratified by source organism. (I) Peer predictors before versus after close-homologue de-duplication. (J) TriStageHLA-BIND AUROC/AUPRC before versus after close-homologue deduplication. (K) Pathogen-stratified AUROC before versus after de-duplication (cross-check).

To render the CEDAR evaluation more stringent, we applied a second, finer-grained de-duplication beyond the exact-match filtering above. Excluding every test peptide within edit distance *≤* 2 of any training peptide (Hamming for equal-length; edit distance for length *±*1–2) removes 2,570 (11.4%) close-homologue instances yet lowers overall AUROC by only 0.18 points (0.9423 to 0.9405) and AUPRC by 1.0 point (0.9505 to 0.9402), leaving the model ranking essentially unchanged (Figure 4J). We further stratified the same benchmark by source organism to rule out pathogen-driven inflation: the largest group—human self/tumor antigens, 78% of positives—yields AUROC 0.933, consistent with the overall 0.942 (HCV 0.992, EBV 0.992, Merkel polyomavirus 0.955; HPV/CMV have too few positives for standalone conclusions), and no pathogen subset collapses after this de-duplication (largest drop 0.0045 for human antigens; Figure 4H, K). Applying the same sequence-only de-duplication to the peer predictors confirms that this robustness is a property of the benchmark rather than of our model: after removing the identical close-homologue subset, the AUROC of MixMHCpred (0.9303*→* 0.9369), NetMHC-pan (0.9239 *→*0.9296), MHCflurry (0.9257*→* 0.9291), BigMHC (0.9195*→* 0.9240), and TransPHLA (0.8934 *→* 0.8970) each remains stable or improves slightly, while TriStageHLA-BIND alone shows a negligible decrease (0.9423 *→* 0.9405; Figure 4I). Together, these two de-duplication levels—exact-match and close-homologue—confirm that our external generalization is robust to both identical and near-identical training examples.

#### 2.3.2 Extrapolation to alleles with limited similarity (distance-based evaluation)

We further examined generalization to alleles that are dissimilar from those seen during training. Following MixMHCpred3.0, we define a distance between two HLA class I alleles based on their 34-residue binding-site pseudo-sequences. Let *a* = (*a*_1_, …, *a*_*J*_) and *b* = (*b*_1_, …, *b*_*J*_) denote the binding-site amino acids (pseudo-sequences) for alleles *a* and *b*, respectively, with *J* = 34. Using BLOSUM62 substitution scores, we compute:

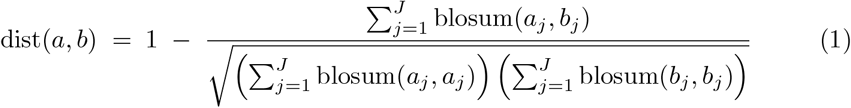

where blosum(*·, ·*) is the BLOSUM62 substitution score. The numerator aggregates cross-allelic similarity across binding-site positions, and the denominator normalizes by allele-specific self-similarity. The resulting value lies in [0, 1], with larger values indicating greater divergence in binding-site biochemistry.

Let *A*_train_ denote the set of HLA alleles present in the training data; for any allele *a*, min 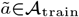 dist(*a, ã*) is the distance from *a* to its most similar training allele under the previously defined dist(*·, ·*). We call an allele “distant” if min 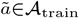 dist(*a, ã*) *>* 0.1. Using this metric, we identified alleles with min 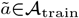 (*a, ã*) *>* 0.1 as relatively distant from the training set. Across all assessed alleles (*n* = 139 total), 21 met this criterion (15.1%), indicating a non-trivial tail of out-of-distribution (OOD) bindingsite compositions (distribution summarized in Fig. 4F). We visualized pseudo-sequence logos for (i) all alleles used in training (Fig. 4D) and (ii) the distant subset (Fig. 4E), highlighting position-wise shifts at canonical pockets.

To quantify extrapolation, we retrained models using only alleles with min 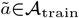 dist(*a, ã*) *≤* 0.1 and then evaluated on the held-out distant set (*n* = 21 alleles; 15.1% of 139). Under this split, the OOD test set comprised 66,870 peptide–allele pairs (25,065 positives; 41,805 negatives). The remaining data were partitioned 9:1 into training (869,048 samples) and validation (96,567 samples). The radar chart in Fig. 4G summarizes performance across metrics: accuracy was 76.39%, recall 75.25%, specificity 77.06%, precision 66.30%, MCC 0.5122, and AUROC 83.66% on this OOD cohort, with precision/recall and MCC remaining stable across peptide lengths—indicating preserved discrimination under distribution shift and suggesting practical extensibility to previously unseen alleles with altered binding-site biochemistry.

Overall, TriStageHLA-BIND generalizes to fully external HLA–peptide pairs assembled independently from the development data and sustains competitive length-wise performance, while retaining measurable accuracy for alleles with limited binding-site similarity to the training distribution.

### 2.4 Ablation studies: overall effects and representation analyses

#### 2.4.1 Overall ablations on AUROC/AUPRC across alleles and lengths

We conducted ablation experiments to quantify the contribution of major components: the convolutional feature extractor (CNN), the mutual-information–based Transductive Information Maximization (TIM) [43] loss, and the initial transfer-learning stage. Figures 5A–C report AUROC and AUPRC aggregated across alleles and peptide lengths, together with length- and allele-stratified summaries.

**Fig. 5.**
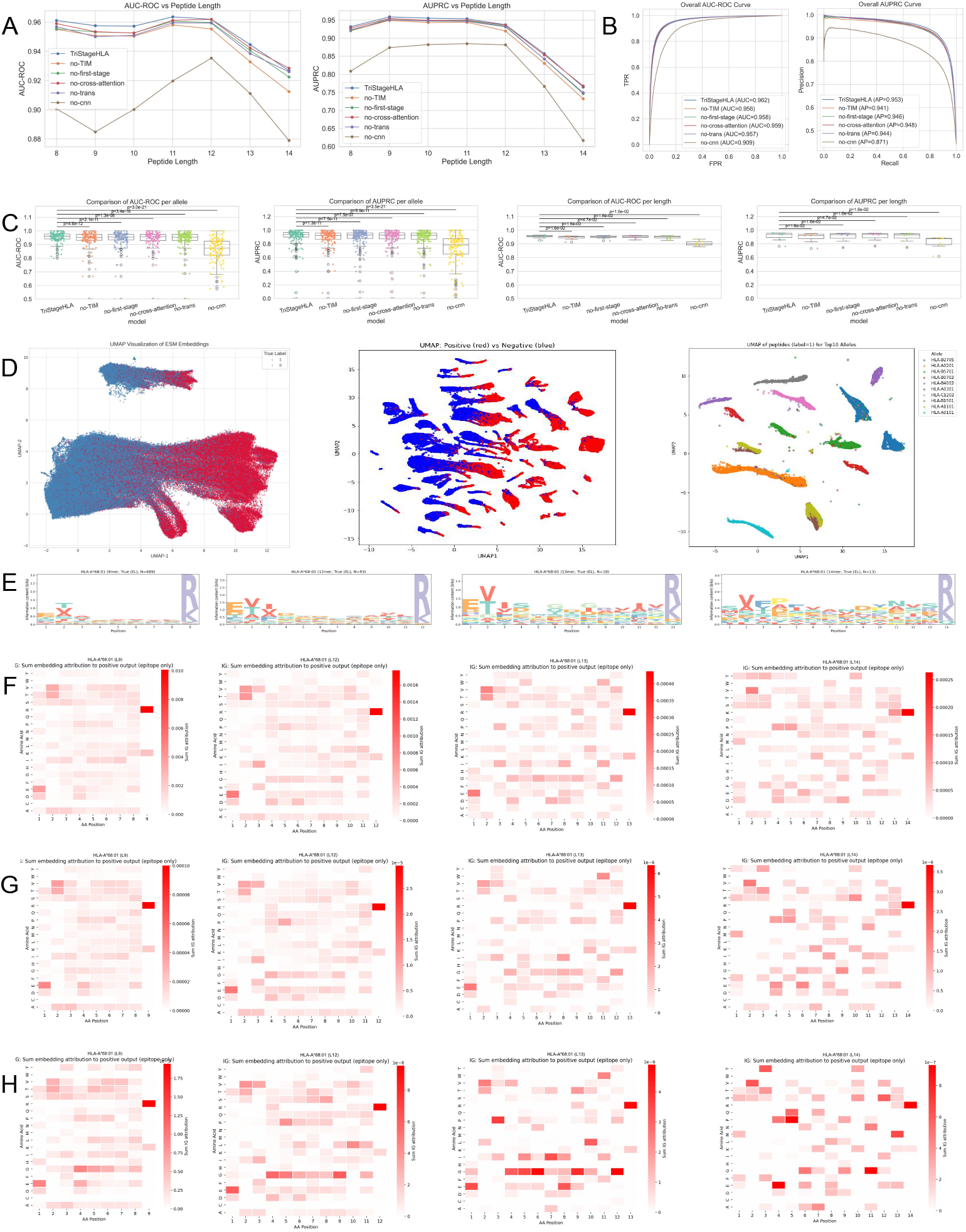
Ablation studies, representation structure, and component-wise interpretability. (A) AUROC/AUPRC across stepwise ablations (−TIM, −cross-attention [44], −DPCNN [46], −LoRA, −frozen encoder) with 95% CIs. (B) Aggregate AUROC with relative drops annotated. (C) Box- and-scatter stratified by allele and length with paired tests versus the full model (BH-adjusted). (D) UMAP embeddings: left, presentation positives vs. negatives; middle, after binding-task training; right, allele-colored clusters for the top-10 alleles by sample size. (E) Length-specific sequence logos (9/12/13/14-mers) for one allele, showing motif shifts. (F) Side-by-side attribution maps across model variants, aligning module contributions. (G) Removing peptide-first transfer learning (directly fine-tuning the *∼* 35M encoder) degrades performance. (H) Removing the TIM objective (CE-only) reduces calibration and AUPRC under imbalance.

Removing the CNN led to a pronounced degradation in overall discrimination, indicating that local motif extraction remains critical for peptide–HLA modeling beyond purely token- or attention-based encodings [44]. Concretely, across different lengths, average AUROC dropped from 95.33% (full) to 90.44% (no-CNN), and average AUPRC from 90.85% to 81.65%. The decline was especially marked at longer peptides; for length 14, AUROC decreased from 92.67% to 87.90% and AUPRC from 76.49% to 61.67%.

In the absence of the TIM loss, overall metrics remained competitive; however, both AUROC and AUPRC deteriorated steeply with the increases in peptide length, suggesting reduced robustness to length-dependent distributional shifts. For example, mean AUROC fell from 95.33% to 94.50% and mean AUPRC from 90.85% to 89.14% across lengths, with larger gaps at 10–12-mers (e.g., length 12 AUPRC 93.71% → 91.98%). Dispersion also increased (see Figure 5C).

Eliminating the initial transfer-learning stage (no-first-stage; i.e., BIND trained from randomly initialized LoRA adapters without the peptide-only PRE pretraining) reduced overall performance by a noticeable margin, consistent with weaker peptide-level priors: across lengths, mean AUROC decreased to 94.92% and mean AUPRC to 89.92%. Separately, removing the Transformer branch of BIND while retaining the PRE pretraining (no-trans; the CNN local-motif branch is kept) produced a comparable but independent drop (mean AUROC 94.87%, AUPRC 89.87%), indicating that global-context modeling adds a small yet consistent increment beyond the local CNN features. The effect was amplified at longer lengths (e.g., length 14 AUPRC 74.67% and 74.92% vs. 76.49% full).

In addition to average performance, dispersion and stability changed across ablations. Both the no-TIM and no-first-stage variants exhibited more extreme values (outliers) in the length- and allele-wise distributions relative to the full model, as visible in Figure 5C. Compared to the complete TriStageHLA-BIND, these ablated variants were less stable across alleles and lengths, particularly at longer peptides. Allele-wise, the full model retained higher central tendency with similar or lower variability (e.g., AUROC mean 94.56% with std 7.66%), while no-CNN showed the largest spread and the lowest mean (AUROC mean 86.27%, AUPRC mean 73.95%), underscoring the primacy of local motif features.

#### 2.4.2 Representation analyses (UMAP and allele-resolved patterns)

To probe how components shape learned representations, we visualized UMAP (Uniform Manifold Approximation and Projection) [45] embeddings from the penultimate layer under different training regimes (Figure 5D). The left panel shows embeddings from the peptide-only pretraining stage: representations are largely compact without clear allele-wise structure. After peptide–HLA binding/presentation training (middle panel), embeddings become structured by allele, with peptide clusters reflecting allele-specific organization rather than a single indistinct mass. The right panel isolates the top-10 most frequent alleles, revealing distinct allele-resolved manifolds; within each, positive and negative peptides remain separable, indicating that the model encodes both allele specificity and class-discriminative signals. These observations support that transfer learning provides a useful initialization for peptide features and that subsequent binding/presentation training introduces allele-aware differentiation while preserving class separation. To further dissect the roles of TIM loss and transfer learning, we examined an allele-resolved case study (HLA-A*68:01). Figure 5E shows sequence logos for 9-, 12-, 13-, and 14-mers; Figures 5F–H provide attribution maps for TriStageHLA-BIND (full), the model without the initial pretraining stage, and the model without TIM loss, respectively.

The complete model aligns more closely with the sequence logos across lengths. For example, at length 13, position 2 shows characteristic preferences (e.g., Valine and Threonine) that are captured more distinctly by the full model than by the no- first-stage variant, which exhibits attenuated sensitivity. The no-TIM variant diverges further: at length 14, position 11 shows an unwarranted emphasis on Glycine, indicating less smooth, allele-consistent preferences. Overall, the full model captures (i) global peptide-level determinants related to epitope potential and (ii) allele-specific constraints; the TIM loss encourages information sharing between peptide and allele embeddings that stabilizes these signals across lengths and helps in data-imbalanced regimes (e.g., longer peptides). Consequently, TriStageHLA-BIND exhibits improved alignment with biochemically plausible motifs and more consistent attribution patterns, particularly at longer lengths where baseline models tend to be less stable.

### 2.5 End-to-end proteome-scale screening

To test the central end-to-end claim—that the staged pipeline yields a more useful candidate list on a realistic proteome-scale screen—we simulated a screen over 300,000 background peptides drawn from the human proteome (absent from all training corpora) together with 14,490 true candidate peptide–allele pairs, and passed them through PRE and then BIND (Figure 6). The peptide-only PRE stage retains 82.4% of true candidates (10,747/13,044) while discarding 87% of background peptides (38,805/299,996 kept); the allele-aware BIND stage then retains 74.4% of true candidate pairs (10,779/14,490) while removing 90% of background pairs. Because the two stages act on different axes—PRE prunes peptides, BIND assigns the correct allele—the pipeline narrows the search without a recall collapse, and peptide-level precision among the top-ranked PRE predictions is high (72.0% in the top 100, 65.8% in the top 1,000, an enrichment of *∼* 17-fold over the 4.2% base rate). Against a no-PRE baseline in which every one of the 300,000 background peptides is expanded across all 139 alleles into 41.7 million pairs, the staged pipeline reduces the computational input roughly 7-fold; measured in terms of the (peptide, allele) pairs that would actually be sent to experimental validation, the saving at matched recall is approximately 2-fold (0.556 vs. 1.144 million pairs at 74.4% recall). PRE’s threshold is a continuous, tunable dial rather than a fixed gate: at higher stringency it discards more background at a bounded recall cost, allowing users to trade validation budget against missed-epitope risk, and the need for a lightweight pre-filter grows precisely because BIND is more accurate and hence more expensive to run at full proteome scale.

**Fig. 6.**
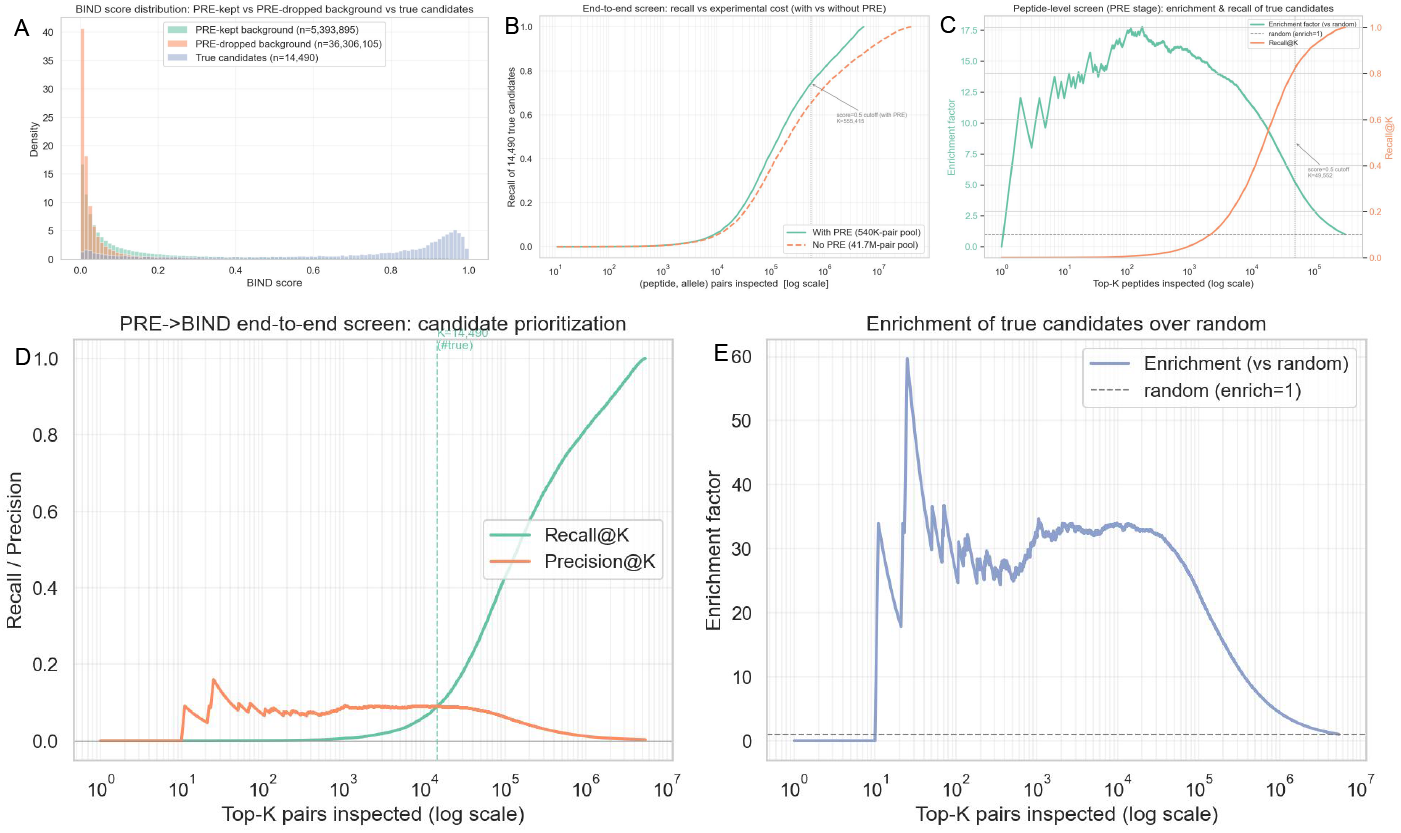
End-to-end proteome-scale screening (PRE*→* BIND) over a proteome-scale candidate pool. (A) BIND score distributions of background peptides retained by PRE versus those removed by PRE, together with the true candidate peptides; the PRE-dropped peptides are also scored low by BIND (only 1.6% receive a BIND score *≥* 0.5), confirming that the two stages agree on what to discard. Recall of the 14,490 true candidate (peptide, allele) pairs as a function of the pairs inspected, comparing the pipeline with PRE (5.4M-pair pool) against a no-PRE baseline (41.7M-pair pool); at matched recall PRE halves the pairs sent to experimental validation (0.56M vs. 1.14M at 74.4% recall). (C) Peptide-level PRE screen: enrichment factor and recall of the 13,044 true candidate peptides ranked among 313,040 pooled peptides (4.2% base rate); the top-100 and top-1,000 recover true candidates at 72.0% and 65.8% precision (*∼* 17-fold enrichment), and a 0.5 score cutoff retains 82.4% of true candidates while removing 87% of background. (D) Pair-level BIND screen: recall@K and precision@K of the 14,490 true candidate pairs among 5.4M scored pairs (peak precision *∼* 9%; a 0.5 cutoff retains 74.4% of true candidate pairs). (E) Pair-level enrichment over random (peak *∼*34-fold).

### 2.6 Immunogenicity prediction

To evaluate T-cell immunogenicity beyond binding alone and to quantify generalization to independent cohorts, we curated positives only from experimentally validated immunogenic peptide–HLA pairs and constructed matched non-immunogenic controls under patient- and study-disjoint splits, so that held-out test cohorts provide clear, reproducible evidence of broad immunogenicity rather than assay- or cohort-specific artifacts; the specific curation, filtering, and partitioning procedures are detailed later in the Methods (Datasets) section. On the held-out immunogenicity test, TriStageHLA-IM achieved an AUROC of 81.4% and an AUPRC of 73.8% (Figure 7). Relative to the strongest immunogenicity-specific baselines, TriStageHLA-IM improves AUROC from 70.7% (ImmuneApp) and 60.1% (BigMHC-IM) to 81.4%, and AUPRC from 63.6% and 51.5% to 73.8%; the modular protein-language-model immunogenicity predictor ImmGenX and the IEDB Class I immunogenicity tool reach AUROC 51.5% and 52.4% (AUPRC 43.1% and 41.4%), respectively, on this benchmark. These results indicate that TriStageHLA-IM separates immunogenic from non-immunogenic peptides with substantially higher discrimination than both immunogenicity-trained and presentation-only methods. Baselines that focus on presentation (e.g., EL/BA-trained models), shown as a sanity-check panel, can rank binders but provide limited incremental signal for immunogenicity on this benchmark. The immunogenicity-trained predictors (ImmuneApp, BigMHC-IM, ImmGenX, and the IEDB immunogenicity tool) perform better than or comparable to presentation-only models, consistent with their immunogenicity supervision, yet each remains below TriStageHLA-IM. Conversely, approaches that augment immunogenicity supervision with large volumes of binding-only data (e.g., PRIME 2.1 variants) may underperform on immunogenicity if binding signals dominate the learned representation; in our evaluation, this led to lower precision–recall behavior compared to TriStageHLA-IM. To verify that this ranking reflects the benchmark rather than an error in how PRIME was invoked, we reproduced the PRIME-versus-presentation comparison on the public BigMHC immunogenicity benchmark (*n* = 937): under the authors’ headline metric (mean PPV_*n*_), PRIME 2.1 (0.2405) scores below both MixMHCpred-3.0 (0.2634) and NetMHCpan-4.1 (0.2551), confirming that PRIME does not, by construction, out-rank presentation-only tools on immunogenicity. The mechanism is specific to our benchmark’s negative construction: because non-immunogenic controls were themselves assayed on the expectation that they bind, the binding signal carries almost no discrimination here (MixMHCpred %Rank AUROC 0.461, NetMHCpan EL-rank 0.508), and PRIME, whose score faithfully tracks MixMHCpred %Rank (Spearman *r* = *−* 0.867), inherits this non-discrimination (PRIME AUROC 0.432). This is an expected property of a clinically realistic benchmark in which both classes pass a binding pre-screen, not a defect in our evaluation pipeline. The binding-rank distribution, the reproduced PPV_*n*_ ranking, and the PRIME-versus-binding correlation are visualized in Supplementary Figure 11.

**Fig. 7.**
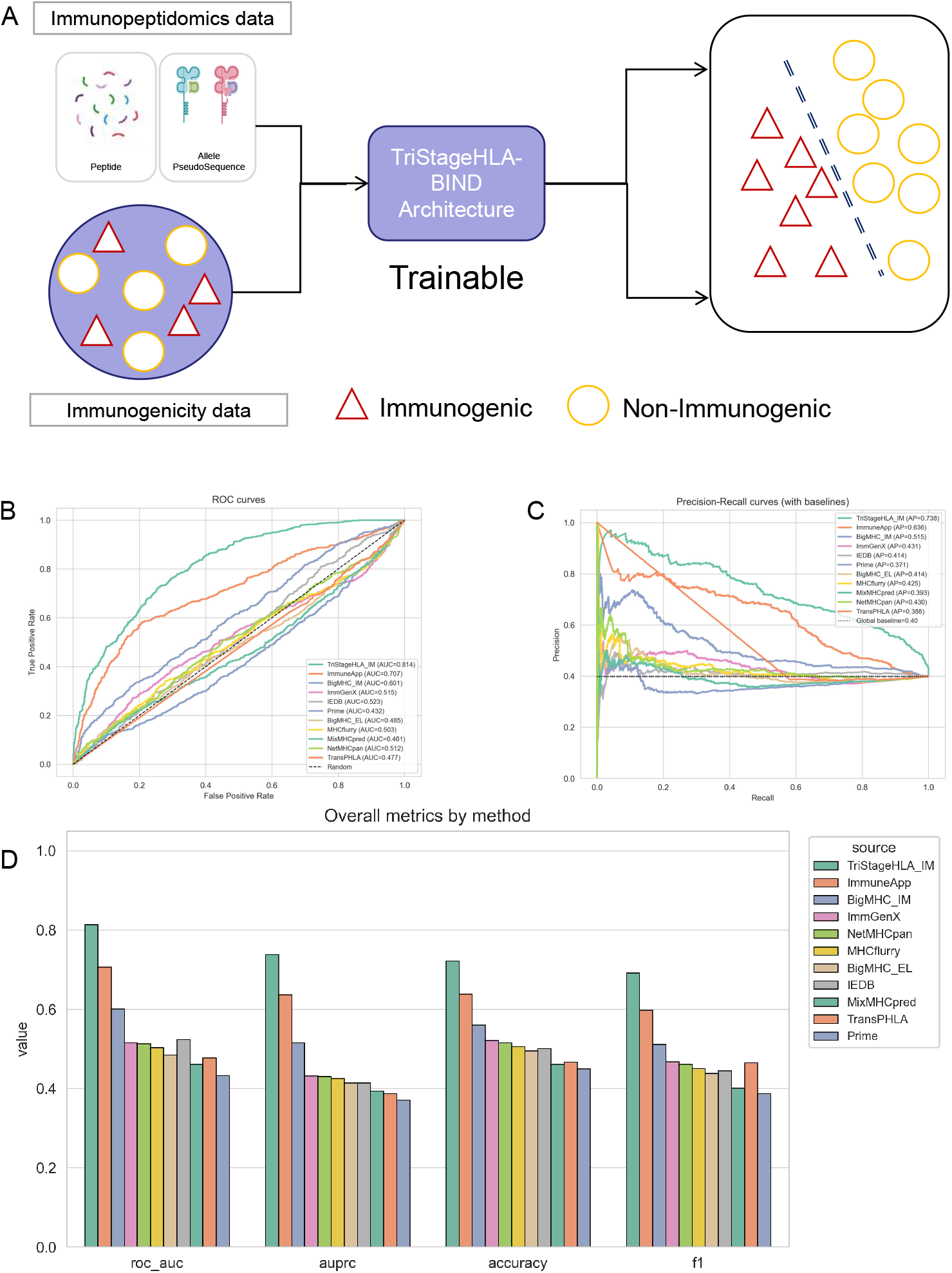
Immunogenicity (IM) prediction on independent tests with baseline comparisons. (A) Schematic overview of the TriStageHLA-IM workflow. (B) AUROC curves and summary AUROC. (C) Precision–recall curves and summary AUPRC. (D) Bar plots of AUROC/AUPRC across the 11 methods—immunogenicity-trained tools together with binding/presentation-only tools, the latter shown as a sanity-check panel; metrics computed from raw logits.

Overall, these analyses support that TriStageHLA-IM captures immunogenic determinants beyond binding, yielding improved AUROC/AUPRC and superior precision at clinically relevant top-*n* thresholds compared with presentation-focused predictors and immunogenicity models trained with broader binding supervision.

## 3 Discussion

Recent advances in LC–MS/MS immunopeptidomics and expanded binding/elution corpora have enabled increasingly precise modeling of HLA class I antigen presentation. Building on these resources, TriStageHLA contributes a compact and parameter-efficient framework that separates peptide-first enrichment, allele-aware binding/presentation, and immunogenicity into coordinated stages. This design preserves end-to-end tractability while improving discrimination across peptide lengths and alleles. From a clinical standpoint, the immediate consequence is a screening work-flow that can run on modest hardware, at proteome scale, and deliver a shortlist of candidates worth experimental follow-up—the step that currently dominates the cost of epitope-based immunotherapy development.

Noticeably, we observe several insights into HLA class I modeling. Firstly, freezing the backbone with lightweight LoRA adapters yields task-specific gains with markedly fewer trainable parameters, enabling deployment at proteome scale without sacrificing accuracy. Second, principled negative construction and strict, non-overlapping data partitions—combined with an external CEDAR assessment and a distance-based OOD protocol—support calibrated evaluation and more robustness estimates. Empirically, TriStageHLA-BIND matches or exceeds strong baselines (MixMHCpred3.0, NetMHCpan4.1b, BigMHC) on AUROC/AUPRC with stable length-wise behavior, while TriStageHLA-IM improves ROC/PR performance on held-out immunogenicity data under calibrated thresholds.

Regarding the modest absolute gains on binding/presentation, we emphasize that established predictors have largely converged on well-studied alleles (all within roughly 93–96% AUROC on combined BA/EL), so raw binding discrimination is near saturation and is not where the remaining headroom lies. TriStageHLA-BIND’s contribution is therefore not a marginal AUROC improvement but parameter-efficient, calibrated, proteome-scale-deployable binding prediction that concentrates its gains on rare and distant alleles where existing tools underperform. The paper’s central claim—that a staged, decoy-light workflow yields a shorter and more immunogenicity-enriched candidate list—is carried by the immunogenicity stage, where TriStageHLA-IM improves AUROC from 60.1% to 81.4% over the strongest immunogenicity baseline, together with the end-to-end screening results showing that the PRE and BIND stages jointly narrow a proteome-scale search by roughly 7-fold at the computational level and approximately 2-fold at the experimental-validation level.

The ablation study clarifies why each component matters. Removing the local-motif branch consistently reduces discrimination, indicating that the model relies on recognizing short sequence patterns as much as on long-range context. Removing the information-sharing constraint that keeps the peptide and HLA views mutually consistent makes predictions less stable, especially for longer peptides and less common alleles. Removing the early, peptide-focused training step lowers accuracy and increases variability, showing that first learning to recognize epitope-like features from abundant peptide data helps even when the final task is allele-specific. The learned representations corroborate this picture: as training proceeds, the model’s internal organization shifts from grouping peptides by sequence alone to separating them by HLA allele and by binding outcome, and the importance it assigns aligns with known peptide-HLA contact regions.

Generalization was evaluated along two axes that matter for deployment. On CEDAR, TriStageHLA-BIND maintains strong AUROC/AUPRC with fewer outliers than presentation-focused baselines, indicating consistent performance across sources. Under binding-site distance stress, retraining on close alleles and testing on distant ones preserves measurable accuracy and AUROC, supporting practical extensibility to under-represented alleles and motivating distance-aware evaluation.

From a high-throughput screening perspective, high precision among top-ranked candidates is critical. The decoupled design of TriStageHLA—with separate LoRA heads for BIND and IM—allows each module to focus on its respective supervision signal without interference. On the immunogenicity module, strict curation of human T-cell assay labels and re-initialized adapters improved performance over binding-trained baselines, suggesting that task-specific fine-tuning beyond binding is beneficial. More broadly, this modular separation clarifies the distinct supervision each stage can reliably leverage.

Limitations remain. Our present focus is HLA-I; extending to HLA-II will require adapted encodings, peptide-length handling, and negative design. Immunogenicity labels used in this study are operationally defined by positive T-cell assay readouts in IEDB; they reflect assay-detectable reactivity and may be influenced by cohort-specific TCR repertoires and prior exposure, rather than an intrinsic peptide property. Regarding external validation, while we excluded exact peptide–allele pairs and over-lapping peptide sequences between CEDAR and IEDB, we cannot fully rule out the effect of close-homologue peptides (e.g., single-residue variants) that may share similar binding motifs and could partially inflate apparent generalization performance. Additional multi-center datasets would strengthen external validity. Although our negative construction and calibration choices aim to reduce bias, remaining batch effects and prevalence shifts can affect thresholds.

Future work will prioritize (i) HLA-II extensions with class-specific pseudosequences and length modeling, (ii) deployment-oriented calibration and thresholding with decision-focused metrics (e.g., top-k hit rate and precision at clinically relevant operating points), (iii) broader external validations spanning multiple laboratories and acquisition protocols, with stricter sequence-similarity filtering to mitigate close-homologue leakage and better assess true out-of-distribution generalization, and (iv) semi-supervised or contrastive objectives to better leverage unlabeled ligandomes, including rare alleles and longer peptides, and (v) prospective validation of end-to-end candidate lists in patient-derived cohorts, which would directly test whether the screening pipeline shortens the path from sequencing data to a therapeutically actionable epitope set. We also plan distance-aware model selection to improve OOD robustness.

In summary, TriStageHLA shows that a compact, transfer-aware and information-theory–aware architecture can deliver strong and robust performance for peptide-first screening and allele-aware binding/presentation, with an additional module for immunogenicity prediction that improves discrimination on curated T-cell assay labels. By aligning sub-tasks, enforcing strict partitions, and stress-testing generalization both externally and by allele distance, TriStageHLA improves decision quality at proteome scale and offers complementary strengths in parameter efficiency, robustness, and deployment readiness. Taken together, the framework positions itself as a practical triage step in the epitope-discovery pipeline—narrowing a proteome-scale search to a short, interpretable, and experimentally tractable candidate set for vaccine and cell-therapy development.

## 4 Methods

### 4.1 Datasets

#### 4.1.1 Allele-aware binding/presentation (BA/EL)

We aggregated peptide–HLA pairs from IEDB ligands (MHC class I) for both binding affinity (BA) and eluted ligand (EL) data (IEDB accessed: 2024–06–06; the snapshot is static and not automatically recompiled). These sources contribute both positive ligands (EL) and quantitative binding affinity (BA). Positives are defined as follows: for BA, peptide–HLA pairs with reported affinity and IC50 ¡ 500 nM are labeled positive (a total of 215,208 BA-annotated pairs); for EL, MS-eluted peptide–HLA pairs in IEDB are labeled positive. Negatives are constructed by randomly sampling peptide fragments from protein sequences and pairing them with alleles. We sample random decoys rather than reuse IEDB’s experimentally confirmed non-binders (a relatively small, allele-biased subset concentrated on a few well-studied alleles), because random proteome sampling yields the broad allele coverage and per-allele 1:1 balance required for unbiased calibration; such experimental negatives are better reserved as independent validation evidence where available. For peptides reported to bind multiple alleles, we treat each distinct (peptide, allele) pair as a separate instance. This allows the model to learn from allele-discordant cases—where the same peptide binds one allele but not another—while retaining promiscuous binders that are biologically relevant, rather than excluding or downweighting them. To maintain approximately 1:1 balance per allele, if an allele’s current negative count already meets or exceeds its positive count, we do not inject additional negatives; otherwise, we inject negatives up to parity with the positive count. Applying these rules yields 1,032,486 peptide–HLA instances for training, including 459,989 ligands (positives) (BA + EL) and 572,494 negatives, with the same per-allele balancing protocol carried over to validation and test. Allele coverage is broad (139 alleles); detailed statistics are provided in the Supplementary Materials.

#### 4.1.2 External generalization set (CEDAR)

Since all available IEDB data have been fully allocated to our internal training and validation (including a held-out internal test set), we construct an external CEDAR evaluation set restricted to human, MHC class I, peptide length 8–14, and MS/ligand evidence. We remove any ligand overlapping with our internal train/validation/test (exact peptide–allele pairs) and additionally exclude any peptide sequence that appears anywhere in our training or evaluation corpora used for peptide-only presentation and allele-aware BA/EL modeling. The resulting positive set contains 11,238 peptide–HLA ligands whose peptides are entirely unseen under the above exclusion. For negatives, we sample 11,238 peptide–HLA pairs such that the peptides are like-wise unseen across all internal splits and corpora. This CEDAR-derived set is used exclusively for external testing, providing a benchmark that is completely decoupled from the IEDB-derived training and internal test data.

#### 4.1.3 Proteome-scale screening simulation (end-to-end)

To evaluate the staged PRE *→* BIND pipeline under realistic conditions, we assembled a screening benchmark from two components. The background set consists of 300,000 human-proteome peptide fragments (8–14-mers) verified to be absent from every internal training, validation, and test corpus of all three stages (peptide-only presentation, allele-aware binding, and immunogenicity), so that none overlaps any development data. The true-candidate set comprises 14,490 (peptide, allele) pairs spanning 139 alleles, drawn from our labeled binding and immunogenicity data (3,539 immunogenic positives, 34 immunogenic negatives, and 10,917 binding-only positives; 13,044 unique peptides). For the no-PRE baseline, each of the 300,000 background peptides is expanded across all 139 alleles into 41.7 million (peptide, allele) pairs, the input size against which the staged pipeline is compared.

#### 4.1.4 Immunogenicity Datasets

We curate a high-specificity immunogenicity dataset from IEDB, restricted to linear epitopes, T-cell assays, MHC class I, human, and any disease, with peptide length 8– 14. To ensure informative supervision, we remove instances lacking explicit four-digit HLA alleles, deduplicate exact peptide–allele duplicates while retaining distinct alleles for the same peptide as separate instances, and exclude negative instances without explicit cohort metadata (number of subjects tested and responded) or with fewer than four subjects tested. Final labels follow IEDB T-cell assay outcomes after filtering. The resulting dataset contains 9,883 instances in total (positive = 3,855; negative = 6,028). We construct instance-level train/validation/test splits with no leakage using a 7:1:2 ratio, and report allele coverage in Supplementary Tables (82 alleles) (IEDB accessed: 2025–04–22).

Given that these labels originate from diverse T-cell assays across different cohorts, it is important to clarify their operational meaning. We define an instance as immunogenicity-positive if the original assay reports a measurable response to the peptide–HLA complex under the specified experimental conditions. This operational definition reflects assay-detectable T-cell reactivity rather than an intrinsic, context-independent property of the peptide. Accordingly, TriStageHLA-IM is trained to predict the probability of a positive T-cell readout in IEDB-style assays, a signal influenced by both peptide characteristics and cohort-specific factors such as TCR repertoire and prior exposure.

### 4.2 Training Overview

We adopt a concise two-stage transfer learning scheme for peptide–HLA class I prediction, plus an independent fine-tuning track for immunogenicity. A pretrained protein language model serves as the shared encoder, augmented with lightweight LoRA adapters; downstream, we combine modest intra-sequence context modeling, explicit peptide–HLA coupling, and a compact classifier. Training emphasizes calibration and robustness under class imbalance. We also conduct targeted ablations by removing the local CNN, the transformer blocks, or the cross-attention to assess each component’s marginal utility. Detailed training schedules, hyperparameters, and model configurations are provided in Supplementary Table 5.

### 4.3 Transfer learning with LoRA

In the presentation pre-screening process, we perform allele-agnostic transfer learning by adapting only LoRA adapters on top of a frozen pretrained encoder. We update the low-rank adapter parameters and a lightweight classifier head while keeping all backbone weights fixed, yielding a peptide-level representation that is subsequently reused. Concretely, each fine-tuned linear projection is parameterized as

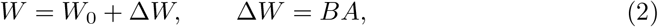

where *W*_0_ denotes the frozen pretrained weight; Δ*W* is the trainable update factorized into a low-rank pair; 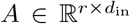 and 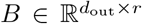 are the learnable LoRA factors with rank *r ≪* min(*d*_in_, *d*_out_); *d*_in_ and *d*_out_ are the input/output channel (feature) dimensions of the projection. For an input 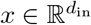, the forward mapping becomes

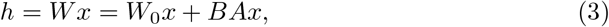

where only *A* and *B* are updated during fine-tuning, ensuring parameter efficiency and stable transfer.

In the allele-aware binding process, we transfer the peptide-level LoRA adapters learned above and continue updating them in a two-branch setting, sharing the same frozen backbone for peptide and HLA inputs but using branch-specific adapters. Only the transferred adapters and a compact classifier are trained, specializing the representation to peptide–HLA interaction while keeping the backbone fixed.

For immunogenicity prediction, we keep the interaction architecture but deliberately do not carry over the adapters from earlier stages. Instead, we attach new LoRA adapters to the same frozen backbone and fine-tune them directly for the immunogenicity objective, decoupling T-cell response learning from biases induced by peptide-only or binding supervision while preserving architectural consistency and parameter efficiency.

### 4.4 Architecture after the frozen backbone

We adopt a dual-branch design for peptides and HLA pseudosequences. The frozen backbone with branch-specific adapters (introduced earlier) produces token embeddings, which are then refined by lightweight Transformer encoders per branch to capture global context. The two streams exchange information via a symmetric cross-attention block that approximates peptide–HLA contact alignment. In parallel, a DPCNN-style residual CNN pyramid per branch extracts local anchor and flanking motifs directly from the original backbone embeddings. The resulting representations are used for downstream classification with a compact head.

#### 4.4.1 Branch-wise Transformer encoders (global context)

Let 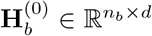 be the backbone token embeddings for branch *b ∈* {*p, h*} (*d*=480). Each branch applies a small stack of encoder blocks to model long-range dependencies. We summarize one block by its essential self-attention and position-wise feed-forward steps:

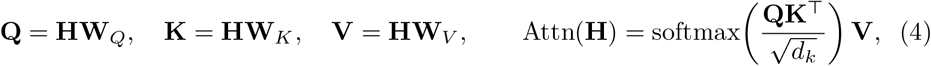

followed by residual connections, layer normalization, and a two-layer feed-forward transformation with nonlinearity (SiLU). Stacking a few such blocks per branch yields refined streams

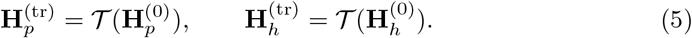

*where* **H** *∈* ℝ^*n×d*^ denotes a generic token-embedding sequence within one branch; 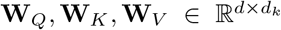 are learned projections to query/key/value spaces; *d*_*k*_ is the attention head dimension; softmax is applied row-wise over token positions; *T* (*·*) denotes stacking encoder blocks (with residual and feed-forward sublayers); *n*_*b*_ is the sequence length of branch *b* (*b*=*p* for peptide, *b*=*h* for HLA).

#### 4.4.2 Symmetric cross-attention (peptide–HLA coupling)

To exchange information across sequences and encourage contact-like alignment, we apply bidirectional cross-attention:

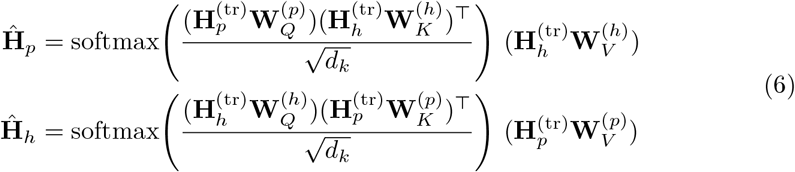

tention map can be interpreted as an approximate peptide–HLA contact alignment signal. *where* 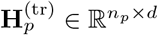 and 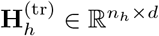 are the branch-wise Transformer-refined token embeddings; 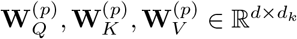 and 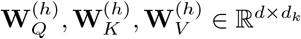 are learned cross-attention projections for peptide and HLA, respectively; the softmax operates row-wise over the attended sequence (e.g., for **Ĥ** _*p*_ it normalizes across *n*_*h*_ HLA positions); 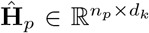 and 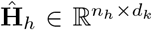 are cross-attended representations (optionally projected back to *d* via an output matrix in implementation).

#### 4.4.3 DPCNN-style residual pyramid (local motif extraction)

Complementing global context, each branch extracts local anchors and flanking motifs from the original backbone embeddings via a DPCNN-style residual pyramid. For branch *b∈ {p, h}* with 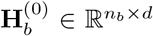, we first form a channel-projected feature map (region embedding) with *c*=256 channels:

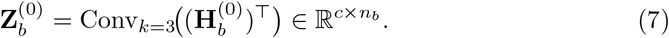

Then we build a residual pyramid by alternating downsampling and small-kernel convolutions:

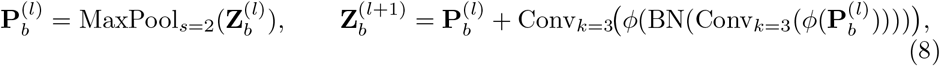

repeating until the temporal dimension is reduced. A final channel-wise batch normalization and temporal pooling yield a local descriptor 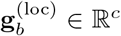 for each branch. *where* (*·*)^*⊤*^ swaps token and channel axes to match 1D-CNN conventions; Conv_*k*=3_ denotes a 1D convolution with kernel size 3 and appropriate padding; *c* is the number of convolutional channels; BN is batch normalization over channels; *ϕ*(*·*) is a point-wise nonlinearity (SiLU); MaxPool_*s*=2_ halves the temporal length with stride 2; index *l* enumerates pyramid levels; Pool_*t*_ (implicit in the last sentence) aggregates over the temporal axis to produce 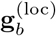.

### 4.5 Training, loss, and optimization

We train TriStageHLA with an information-theoretic objective inspired by TIM, designed for few-shot and imbalanced settings common in HLA data (uneven allele sample sizes and skewed peptide-length distribution, with 9-mers dominating). The objective combines supervised fitting with a mutual-information surrogate that (i) sharpens per-sample predictions and (ii) prevents collapse toward the majority class:

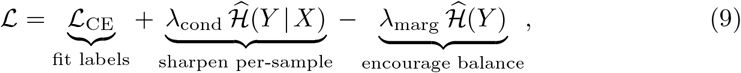

where the supervised cross-entropy is

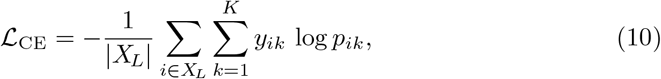

and the empirical entropies are

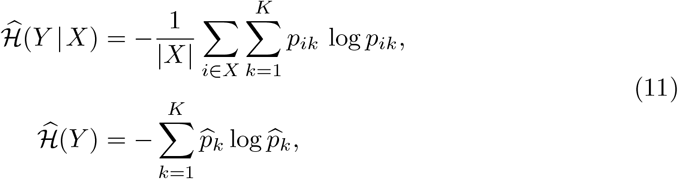

with the empirical label marginal

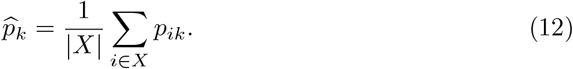

This realizes a batch-wise surrogate of *I*(*X*; *Y*) = *H*(*Y*) *− H*(*Y* | *X*): decreasing 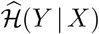 extracts confident signal from scarce alleles and rare peptide lengths, while increasing 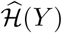 counteracts bias toward dominant classes (e.g., 9-mers).

*where X*_*L*_ is the labeled subset used by the CE term and *X* is the batch used |*X*| are set sizes; *y*_*ik*_ *∈ {*0, 1*}* with ∑_*k*_ *y*_*ik*_ = 1 for labeled data; *p*_*ik*_ are predicted for entropy statistics (may include labeled and weakly/unlabeled samples); |*X*_*L*_| and class probabilities from logits via sigmoid/softmax; *K*=2 (binary); *λ*_cond_, *λ*_marg_ *≥* 0 are tuned on validation.

Optimization details. We use Adam with weight decay and cosine learning-rate decay with warmup (unless otherwise stated), weighted sampling to approximate balanced mini-batches without altering dataset priors, and Xavier initialization for newly introduced linear layers. Early stopping monitors validation AUROC computed on raw logits (no terminal sigmoid/softmax). Checkpoints are saved on AUROC improvement and the best model is restored for evaluation; per-sample probabilities are exported to support downstream calibration.

## 5 Compared methods

### 5.1 Allele-agnostic, peptide-only pretraining

We compare our peptide-only pretraining against a single peptide-centric baseline:

- **TransHLA** is a peptide-only epitope predictor that combines Transformer and residual CNN modules on top of ESM2-derived [47] sequence and structure embeddings to classify whether a peptide is likely to be HLA-presented without requiring allele input. In Step 1, we use TransHLA strictly as the peptide-only reference baseline.

### 5.2 Allele-aware peptide–HLA binding/presentation

We benchmark against five widely used MHC-I predictors covering binding affinity (BA) and/or eluted ligand (EL) tasks:

- **BigMHC-EL** leverages transfer learning from large protein language models and fine-tunes on EL data to produce presentation scores, emphasizing calibrated outputs under class imbalance.
- **MixMHCpred3.0** constructs allele-specific positional weight matrices (PWMs) from mass-spectrometry ligandomes for 8–14-mers, capturing pocket-specific motif preferences for presentation scoring.
- **MHCflurry2.0** outputs BA via an ensemble of two-/three-layer MLPs; an antigen-processing CNN trained with N-/C-terminal flanks augments BA; EL is obtained by logistic regression over BA and processing scores. Its MHC encoding extends NetMHCpan pseudosequences by three residues for finer allele discrimination.
- **NetMHCpan4.1b** jointly models BA and EL using an ensemble of 100 single-layer networks on an inferred peptide 9-mer core and a fixed 34-residue MHC pseudosequence selected by proximity (within 4 °A) to the bound peptide.
- **TransPHLA** [48] encodes peptides and NetMHCpan-style MHC pseudosequences with transformer layers followed by ReLU-activated fully connected projections. To avoid output saturation that can hinder threshold-free metrics (AUROC/AUPRC), we evaluate scores without a terminal softmax.

### 5.3 Immunogenicity

For T-cell immunogenicity, we evaluate immunogenicity-trained predictors together with presentation-oriented baselines from Step 2:

- **BigMHC-IM** adapts BigMHC to immunogenicity labels, fine-tuning contextual embeddings on immunogenic/non-immunogenic outcomes while retaining transfer from presentation tasks.
- **PRIME 2.1** trains directly on immunogenicity to capture TCR-recognition–related signals on top of presentation features, outputting pMHC-triggered response scores.
- **ImmuneApp** [49] predicts peptide immunogenicity alongside eluted-ligand presentation; we use its immunogenicity score for this task.
- **ImmGenX** [50] is a modular protein-language model for CD8^+^ T-cell epitope immunogenicity that encodes peptide–HLA inputs and an optional TCR module; we use its pMHC-only immunogenicity score.
- **IEDB immunogenicity** [51] is the IEDB Class I immunogenicity predictor based on the physicochemical properties of presented peptides; we use its pMHC immunogenicity score directly.
- **BigMHC-EL, MixMHCpred3.0, MHCflurry2.0, NetMHCpan4.1b, TransPHLA**: used as presentation-based comparators on the immunogenicity task for reference; where applicable, we use calibrated or raw scores without additional softmax to maintain AUROC/AUPRC comparability.

### 5.4 Evaluation metrics

We evaluate with accuracy (ACC), area under the ROC curve (AUROC), area under the Precision–Recall curve (AUPRC), and Matthews correlation coefficient (MCC).

***Accuracy (ACC)***.

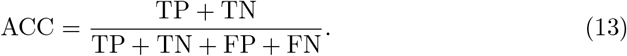

***AUROC***.

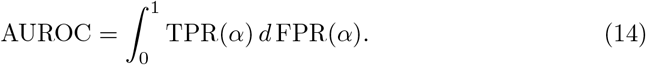

***AUPRC***.

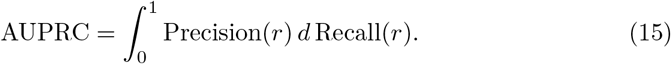

***MCC***.

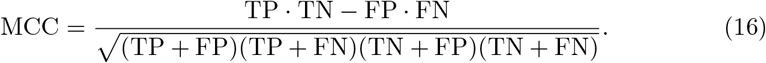

*where* we evaluate binary labels *y*_*i*_ *∈* {0, 1} with continuous model scores *s*_*i*_ *∈* ℝ (no terminal softmax); a decision threshold *τ* induces predictions *ŷ*_*i*_ = ⊮ {*s*_*i*_ *≥ τ*} and confusion-matrix counts TP, TN, FP, FN (default *τ* =0.5 for ACC/MCC; threshold is swept over unique *s*_*i*_ for AUROC/AUPRC). True positive rate TPR 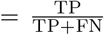 and false positive rate FPR 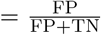 trace the ROC as the threshold *α* varies. Precision 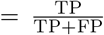 and Recall 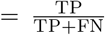 trace the PR curve indexed by *r*. Numerical areas (AUROC/AUPRC) are computed by trapezoidal integration; we report positive-class AUPRC.

## Data Availability

All datasets used in this study are publicly available from the Immune Epitope Database (IEDB), CEDAR, and TransHLA. Inclusion criteria, preprocessing (filtering, deduplication, negative construction), and the train/validation/test splits are described in Materials and Methods. Exact peptide lists, split indices, and metadata (HLA alleles, assay types, labels) are provided in the Supplementary Materials, mirrored in the project repository, and additionally hosted on Google Drive (TriStageHLA data).

## Code Availability

Source code for data preprocessing, feature extraction, model training, evaluation, and figure generation is available at https://github.com/SkywalkerLuke/TriStageHLA under the MIT License. Environment files (conda/requirements), training scripts, and configuration files are included to reproduce all benchmarks.

